# A data-driven model for the assessment of Tuberculosis transmission in evolving demographic structures

**DOI:** 10.1101/112409

**Authors:** Sergio Arregui, María José Iglesias, Sofía Samper, Dessislava Marinova, Carlos Martín, Joaquín Sanz, Yamir Moreno

## Abstract

In the case of tuberculosis (TB), the capabilities of epidemic models to produce quantitatively robust forecasts are limited by multiple hindrances. Among these, understanding the complex relationship between disease epidemiology and populations' age structure has been highlighted as one of the most relevant. TB dynamics depends on age in multiple ways, some of which are traditionally simplified in the literature. That is the case of the heterogeneities in contact intensity among different age-strata that are common to all air-borne diseases, but still typically neglected in the TB case. Furthermore, whilst demographic structures of many countries are rapidly aging, demographic dynamics is pervasively ignored when modeling TB spreading. In this work, we present a TB transmission model that incorporates country-specific demographic prospects and empirical contact data around a data-driven description of TB dynamics. Using our model, we find that the inclusion of demographic dynamics is followed by an increase in the burden levels prospected for the next decades in the areas of the world that are most hit by the disease nowadays. Similarly, we show that considering realistic patterns of contacts among individuals in different age-strata reshapes the transmission patterns reproduced by the models, a result with potential implications for the design of age-focused epidemiological interventions.

**Significance Statement:** Even though tuberculosis (TB) is acknowledged as a strongly age-dependent disease, it remains unclear how TB epidemics would react, in the following decades, to the generalized aging that human populations are experiencing worldwide. This situation is partly caused by the limitations of current transmission models at describing the relationship between demography and TB transmission. Here, we present a data-driven epidemiological model that, unlike previous approaches, explicitly contemplates relevant aspects of the coupling between agestructure and TB dynamics, such as demographic evolution and contact heterogeneities. Using our model, we identify substantial biases in epidemiological forecasts rooted in an inadequate description of these aspects, both at the level of aggregated incidence and mortality rates and their distribution across age-strata.

---

The control of TB is one of the largest endeavors of Public Health authorities ever since the bacterium that causes it –*Mycobacterium tuberculosis*– was discovered (1). Recently, the development of global strategies for diagnosis and treatment optimization has led to TB burden decay worldwide (2), to the point that the End TB Strategy has allowed the scientific community to think that its eradication before 2050 is possible (3, 4). Nonetheless, such goal is yet far away, and TB remains a major Public Health problem (5–7), being responsible for 1.7 million deaths worldwide in 2016 (4). These dramatic data evidence the need of new epidemiological measures and pharmacological resources (8). In the task of forecasting the potential impacts of such new interventions, epidemiological models of TB transmission constitute a fundamental resource to assist decision making by Public Health agents (9).

Among the various limitations that TB modeling has to face in this context, achieving a proper description of the multiple ways whereby TB dynamics couples with populations’ age structure has been pointed out as one of the most critical (5, 10). For example, patients’ age is strongly correlated to the type of disease that they tend to develop more often, as well as to the probability of developing active TB immediately after infection (usually called “fast progression” (8)). This way, while a larger fraction of children younger than 15 years of age develop non-infectious forms of extra-pulmonary TB with respect to adults (25% vs 10% (8, 11, 12)), the risk of fast progression is larger in infants (50% in the first year of life), then decays (20-30% for ages 1-2; 5% for 2-5 and 2% for 5-10), and raises up again in adults (10-20% for individuals older than 10) (13). Additionally, transmission routes of TB, being this a paradigmatic air-borne disease, are expected to show significant variations in intensity across age (14, 15). The empirical characterization of these contact structures constitutes an intense focus of research in data-driven epidemiology of air-borne diseases (16), and their influence on the transmission dynamics of diseases like influenza has been recently explored with relevant implications (17, 18).

Thus, if subjects’ age modifies the disease-associated risks at the level of single individuals, it is likely to expect that changes in the demographic age-structure at the population level will impact TB burden projections. This is mainly due to the slow dynamics that is characteristic to TB, which forces modelers to describe the evolution of the disease during long periods of time, typically spanning several decades. These timescales are rather incompatible with the assumption of constant demographic structures, at least nowadays, since world-wide human populations are presumed to age from the current median of 30 years old to 37 in 2050 (19). And yet, achieving a sensible description of TB transmission able to capture the effects of time-evolving demographic structures remains an elusive goal in TB modeling. Demographic dynamics are traditionally neglected in TB transmission models, the same way that contact structures are assumed to be homogeneous across age-groups (8, 20, 21).

In this work, we incorporate empirical data on demographic dynamics and contact patterns around classical formulations of TB spreading models, thus unlocking less biased descriptions of the spreading dynamics of the disease. To this end, we present a TB spreading model (Figure 1A) whereby we provide a data-driven description of TB transmission that presents two main novel ingredients with respect to previous approaches. First, our model incorporates demographic forecasts by the United Nations (UN) population division (19) (Figure 1B) in order to describe the coupling between demographic evolution and TB dynamics. Secondly, the model integrates region-wise empirical data about age-dependent mixing patterns adapted from survey-based studies conducted in Africa and Asia (22–26) (Figure 1C), instead of assuming that all the individuals in a population interact homogeneously, as traditionally considered in the literature (8, 20, 21).

**Fig. 1.**
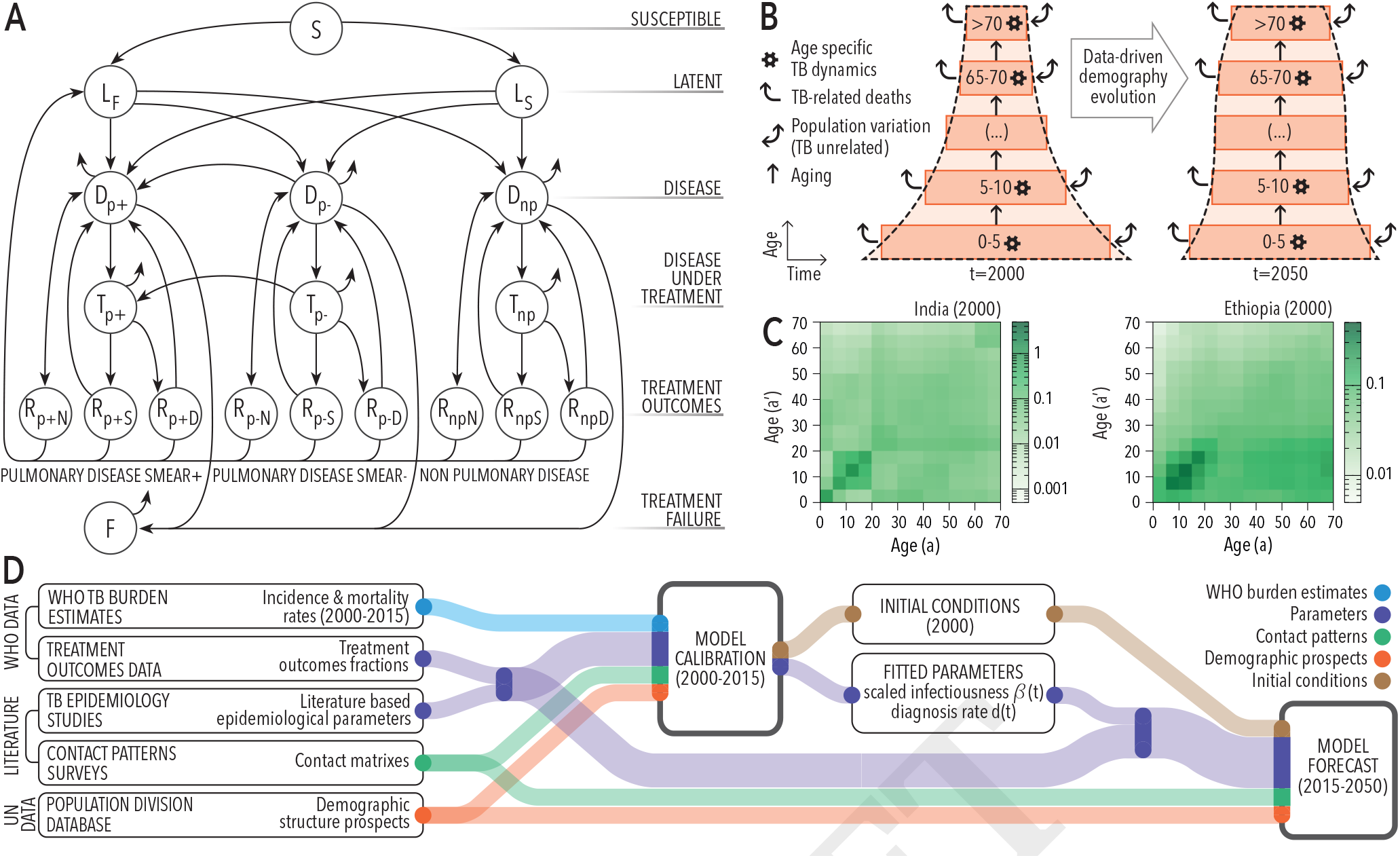
Model description. (A): Natural History scheme of the TB spreading model. S: susceptible. L: latent. D: (untreated) disease, T (treated) disease, R recovered, F: failed recovery. Types of TB considered: p+: Pulmonary Smear-Positive, p-: Pulmonary Smear-Negative, np: Non-pulmonary. Treatment outcomes: *R*_*N*_: Natural recovery, *R*_*S*_: Successful treatment, *R*_*D*_: Default (abandon of treatment), F: treatment failure. (B): Scheme of the coupling between TB dynamics and demographic evolution. The transmission model summarized in panel A describes the evolution of the disease in each age group, including the removal of individuals due to TB-mortality (curved arrows). The evolution of the total volume of each age-strata is corrected (bi-directional arrows: TB unrelated population variations) to make the demographic pyramid evolve according UN prospects. (C): empirical contact patterns used for African and Asian countries. (D): Data flow scheme. Epidemiological parameters, contact matrices, and demographic prospects are used to calibrate the model, with the goal of reproducing observed TB incidence and mortality trends during the period 2000-2015. As a result of model calibration, scaled infectiousness, diagnosis rates and initial conditions of the system in 2000 are inferred. These elements are then used (along with epidemiological parameters, contacts and demographic data) to extend model forecasts up to 2050. For further details regarding model formulation and calibration, the reader is referred to the SI appendix.

Upon model calibration in some of the countries most affected by the disease in 2015 and subsequent simulation of TB transmission dynamics up to 2050 (Figure 1D), we scrutinize the implications derived from integrating these pieces of empirical data within our model, and discuss their impact on the forecasts produced, both at the level of aggregated incidence and mortality rates and on their distributions across age-strata.

## Results

### Baseline forecasts of TB incidence and mortality

To illustrate the ability of our method to reproduce current epidemic trends in different scenarios, the model was applied to describe the TB epidemics in India, Indonesia, Nigeria and Ethiopia (Figure 2). These countries, which accumulate as much as ~40% of the total TB burden world-wide in 2015, were selected because of their different temporal evolution trends, current and prospected demographic profiles and geographic locations. Remarkably, our model does not predict, in general, a sustained decrease in TB burden for the decades to come in these cases, whose incidence rates (per million habitants and year) range between 1246 (524-2124 95% C.I.) (Ethiopia) and 3669 (2348-5247 95% C.I.) (Indonesia), in 2050. Additionally, we extended these analyses to the top-twelve countries suffering from the highest absolute TB burden levels in 2015, producing satisfactory fits in all cases (see supplementary Information (SI), figure S1 and Table S1).

During simulations, our model produced TB detection ratios (TB cases diagnosed divided by new incident cases) that strongly correlate to the notification rates across countries reported by WHO (27) (See SI: figure S2. Pearson correlation r=0.96, p=4.3E-6). Model-based case detection ratios are systematically larger than notification rates, which is an expected result, coherent with the fact that a fraction of all diagnosed cases is not notified to the WHO surveying system.

Regarding confidence intervals, colored areas in Figure 1 quantify the contribution to global uncertainty that stem from the different types of input data processed by the model. This includes epidemiological parameters (purple), demographic data (orange), contact patterns (green), and, most importantly, WHO-burden estimations (blue). Complementarily, in the figure S3 of the SI appendix, the individual contribution of each epidemiological parameter is further disclosed in an exhaustive sensitivity analysis. Of all the different individual sources of uncertainty that are susceptible to impact model’s forecasts, current WHO estimates for TB burden levels is the only one that introduces more than a 15% deviation with respect to central estimates (the uncertainties in total number of TB cases prospected in 2000-2050 that are propagated from WHO-data span from 36% (Ethiopia, lower limit) to 92% (Nigeria, upper limit) with respect to central expectations).

## Effects of populations aging on aggregated TB forecasts

As it can be deduced from the demographic pyramids in Figure S1 of the SI, all countries analyzed in this work are experiencing population aging to some extent, consistent with the overall trend that is forecasted for global human populations during the same period (19). The four countries selected in figure 2 lie at different points of the demographic transition by the beginning of the period under analysis (year 2000), and are expected to evolve at different paces into more or less aged populations by 2050.

**Fig. 2.**
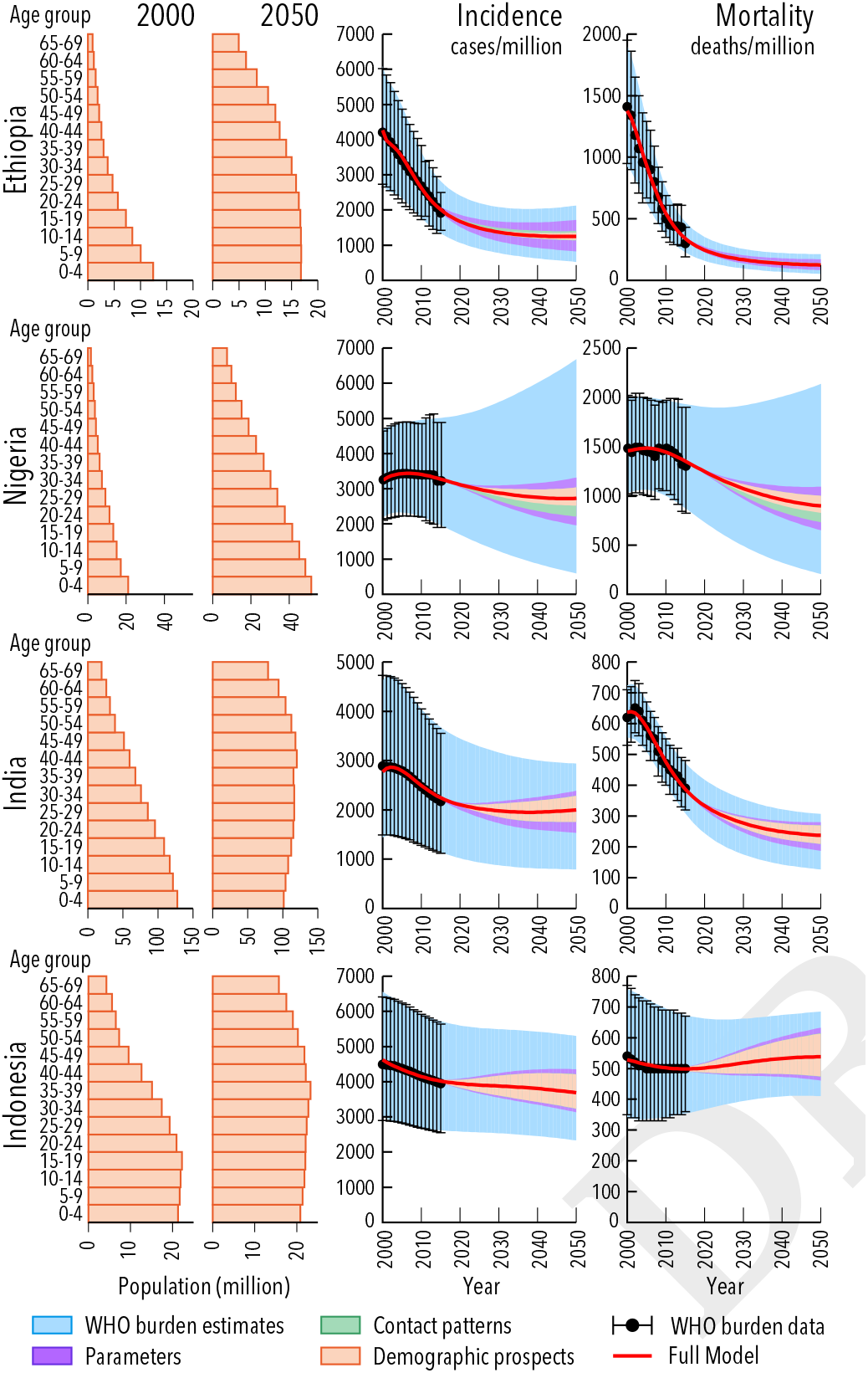
Population structure at 2000 and 2050 (projection), and annual incidence and mortality rates predicted by our model in 2000-2050 for Ethiopia, Nigeria, India and Indonesia. Colored areas represent 95% confidence intervals. The contribution to overall uncertainty stems from each of the four types of input data is disclosed.

In order to isolate the influence of populations aging on model outcomes, we compared our model with a simplified version where demography evolution is neglected as done in previous approaches (8, 20, 21) (reduced model 1). In this reduced model the demographic structures are taken from their initial configuration in 2000 and remain static until 2050. Our results shows that the demographic evolution leads to a systematic and significant increase in the prospected incidence rates, which is variable in size across countries (Figure 3A: relative increase in incidence in 2050: full vs reduced model 1: India: 39.6% (13.9-63.6 95% C.I.), Indonesia 23.4% (7.9-36.5 95% C.I.), Ethiopia 56.0% (29.2-62.1 95% C.I.), Nigeria 34.5 % (9.1-42.9 95% C.I), see also SI appendix, figure S1 and table S2 for equivalent results in other countries). Furthermore, the relative variation between incidence forecasts obtained from the full and the reduced model by 2050 significantly correlates with the intensity of the aging shift, as given by the change in the fraction of adults (age>15 years) in 2000-2050 (Figure 3B, Pearson correlation r=0.66, p=0.02). This is indeed a natural consequence, since adults are burdened with higher incidence rates than children, and thus, populations’ aging implies a relative increase of the demographic strata that is most affected by the disease (adults), in detriment of children, among whom TB incidence is lower (Figure 3C).

**Fig. 3.**
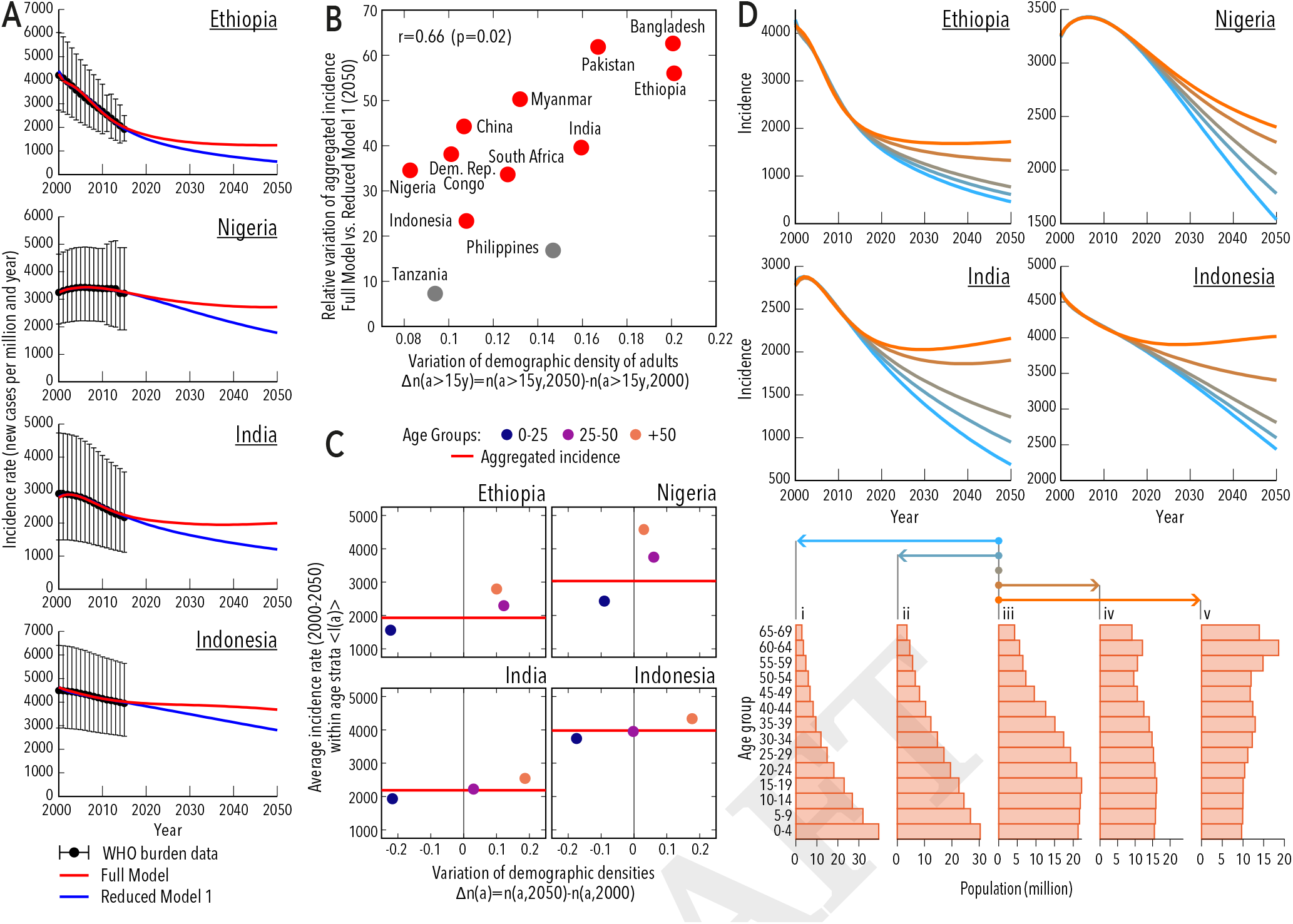
Effects of demographic dynamics on model forecasts. (A): Incidence rates from 2000 to 2050 obtained from the full model (red) and reduced model 1 (constant demography, blue). Relative variation in incidence in 2050: full vs reduced model 1: India: 39.6% (13.9-63.6 95% C.I.), Indonesia 23.4% (7.9-36.6 95% C.I.), Ethiopia 56.0% (29.2-62.1 95% C.I.), Nigeria 34.5 % (9.1-42.9 95% C.I). (B): Relative variation of aggregated incidence at 2050 for the top 12 countries with highest absolute TB burden in 2015 versus variation in the fraction of adults in the population during the period 2000-2050. In all countries but Tanzania and Philippines, in grey in the figure, the variations in incidence are significant at a nominal p=0.05. (C): Age specific average incidence rate of TB vs. variation of age-strata population density in 2000-2050. Older individuals are, at the same time, those affected by higher TB incidence rates and those whose presence in the population is increasing as a results of populations’ aging. (D): Incidence projections for synthetic scenarios of demographic evolution, including transition towards younger populations (iii to i and iii to ii), static populations (iii remaining constant), and realistic transitions representing populations’ aging (from iii to iv and from iii to v). Pivotal demographic structures corresponding to stages i, iii and v are taken from actual examples (Ethiopia in 2000, Indonesia in 2000 and China in 2050), and normalized to a common total population to rule out hypothetical system volume effects. Stages ii and iv are obtained upon linear interpolation. In each panel, the demographic evolution of each country is substituted by these synthetic scenarios: demographic transitions that go from stage iii in 2000 to different ending points in 2050, as indicated by color arrows. Model calibration is repeated in each case.

Next, we built a series of synthetic demographic evolutions to simulate different scenarios (Figure 3D). To this end, we used three pivotal examples extracted from actual cases of populations featuring young, triangular demographic pyramids (figure 3D, stage i, extracted from Ethiopia in 2000), aged, inverted-pyramids (stage v, extracted from China, 2050); as well as intermediate situations (stage iii, extracted from Indonesia, 2000, and stages ii and iv, built upon linear interpolation). Making use of these pivotal populations, we built synthetic transitions among them occurring in the period 2000-2050, which we then integrate in our TB model, in the four countries analyzed, instead of their own real demographic projections. As we can see in figure 3D, population aging appears associated to increased incidence rates, while eventual transitions towards younger populations would cause incidence forecasts to decline faster.

To further validate the general character of these results, we have performed a series of robustness tests in scenarios that go beyond the assumptions made in our modeling framework. That includes comparing full and reduced model 1 under the assumption of highly biased input data (SI, Figure S4), swapping contact structures across continents (SI, Figure S5), interrupting the time-evolution of the fitted parameters after 2015 (see Methods and SI figure S6) as well as dispensing with the independent calibration of the reduced model to rule out the possibility of these differences arising from technical artifacts during model calibration (SI, Figure S7). Remarkably, under all these alternative scenarios, the comparison between full and reduced model remains valid.

Collectively, our results show that ignoring the populations aging within TB spreading models generates forecasts of aggregated burden that are systematically and significantly lower than those obtained when this ingredient is taken into account.

## Effects of aging on age-specific burden levels

Next, we interrogated whether the effect of aging on TB burden estimates is only due to a relative increase of the age-strata more hit by the disease (i.e. adults), or if, in turn, significant increases in the incidence rates within age groups can be identified.

In figure 4, we show, for one example per continent –India and Ethiopia–, the infection matrices between age groups as described by each model, and their difference. The entry (*a*, *a*′) of these matrices represents the prospected number of infections (in 2050) from age-group *a* (infection source) to *a*′ (infection target) per year per million people in group *a*′. For both countries, the differences between full and reduced model 1 point to a systematic under-estimation of the number of infection events caused by adults as a consequence of ignoring demographic dynamics, as well as an over-estimation –only appreciable in India–, of infections caused by children during the period under analysis. Furthermore, once contagions are aggregated across infection sources within each target agegroup (Figure 4, age-specific infection rates histograms, built as column-wise marginal sums of the infection matrices), significant differences between age-specific infection rates arise in both countries, mainly in adult age strata, where the full model predicts systematically larger incidence levels than the reduced model 1.

**Fig. 4.**
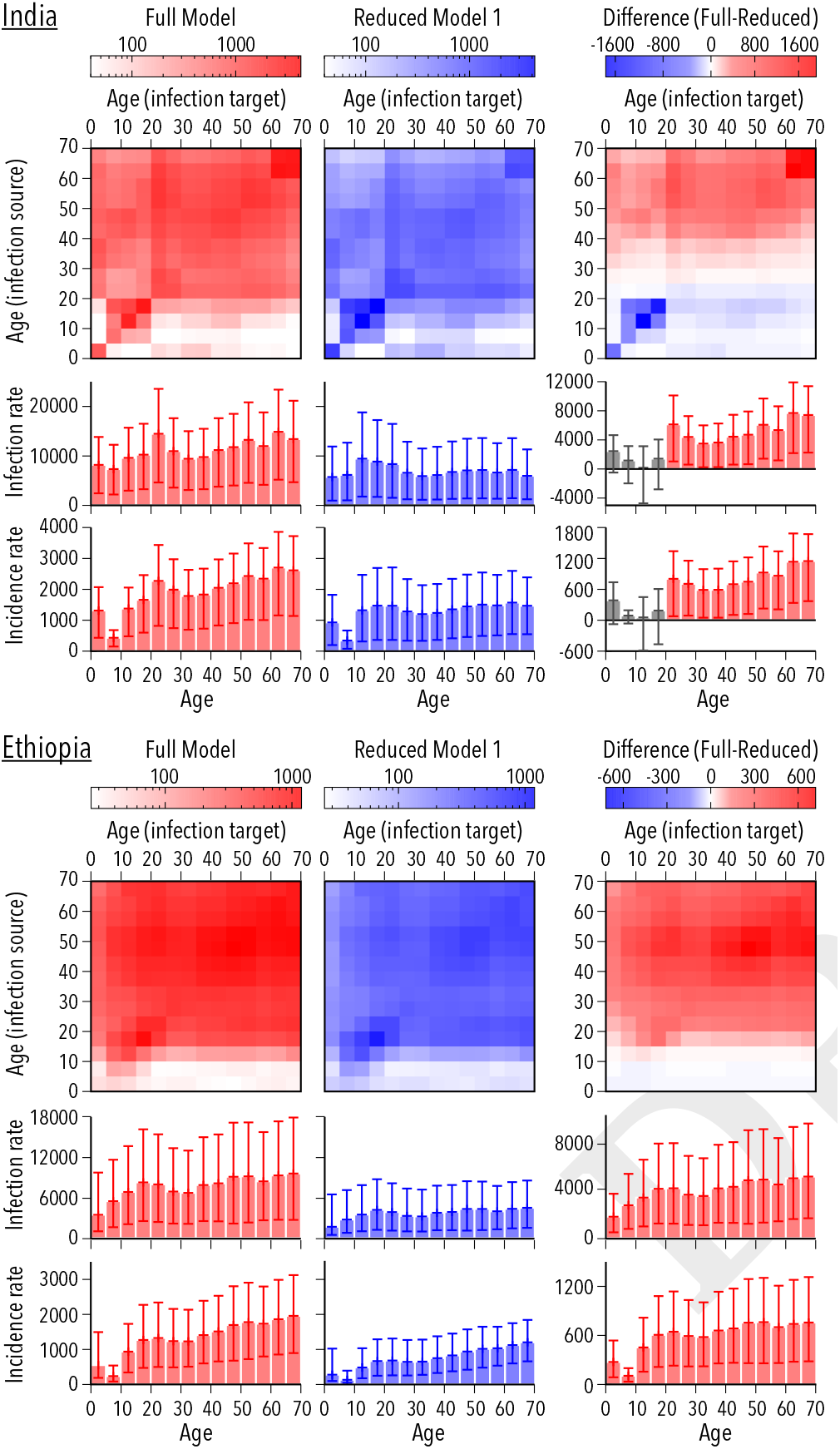
Age-to-age infection rate matrices (number of infections from age group a to age group a’ per year per million people in target age-group a’); and age-specific infection and incidence rates forecasted in 2050 for India and Ethiopia (number of contagions, or new active TB cases, respectively) per year and million individuals in a give age-group). In the left column, the forecasts derive from the full model, and, in the central column, from reduced model 1 (constant demography). In the right column we represent the difference (full - reduced model 1) of these three observables: infection matrices, age-specific infection rates and age-specific incidence rates. Differences in incidence and infection rates are shown in grey when they are not statistically significant. Neglecting demographic dynamics appears associated to an underestimation of infections caused by adults in both countries, and an overestimation of infections from children, mostly in India. At the level of infection/incidence rates, the full model produces larger age-specific infection and incidence rates than the reduced version, more intensely among adults.

This ultimately translates into an increase in age-specific incidence rates of active TB cases (Figure 4, age-specific incidence histograms), which can be easily interpreted attending to the larger probabilities of developing the most infectious forms of pulmonary TB that adults experiment with respect to children (8). Adults, whose proportion increases in the system as a result of considering populations’ aging, not only constitute the part of the demographic pyramid most hit by the disease, but also the one that contributes the most to overall spreading. Therefore, including populations’ aging on model dynamics causes not just an increase in the aggregated burden levels across all age-groups, but also within age-strata.

## Effect of contact patterns heterogeneities

After discussing the impact that demographic dynamics has on model outcomes, we inspected what are the effects of including contact patterns in the TB model forecasts, either at the level of aggregated rates or within age-specific strata. To do so, we have built a second reduced model where the empirical contact matrices estimated from survey studies conducted in Africa and Asia (22–26) are substituted by the classical hypothesis of contacts homogeneity (see Methods and SI section 2.3 for further details).

In figure 5, we represent the infection matrices that derive from the full and the reduced model 2 for India and Ethiopia in 2050. Clearly, empirical contact patterns reshape the distribution of contagions among age-groups, giving a larger importance to assortative infections that take place among individuals of similar ages –specially between adolescents and young adults–, while penalizing infections from children to adults, or vice-versa. As a result, in this case, the infection and incidence rates of TB among children are higher in the reduced model, while the full model predicts more infection and disease burden among adults, with slight variations between the two countries that are due to the different contact data used in each case in the full model. In all the countries analyzed, the opposite directions of the differences between full and reduced model that are found in children versus adults tend to compensate each other. This results into similar global incidence rates produced by both models (see SI appendix, figure S8).

**Fig. 5.**
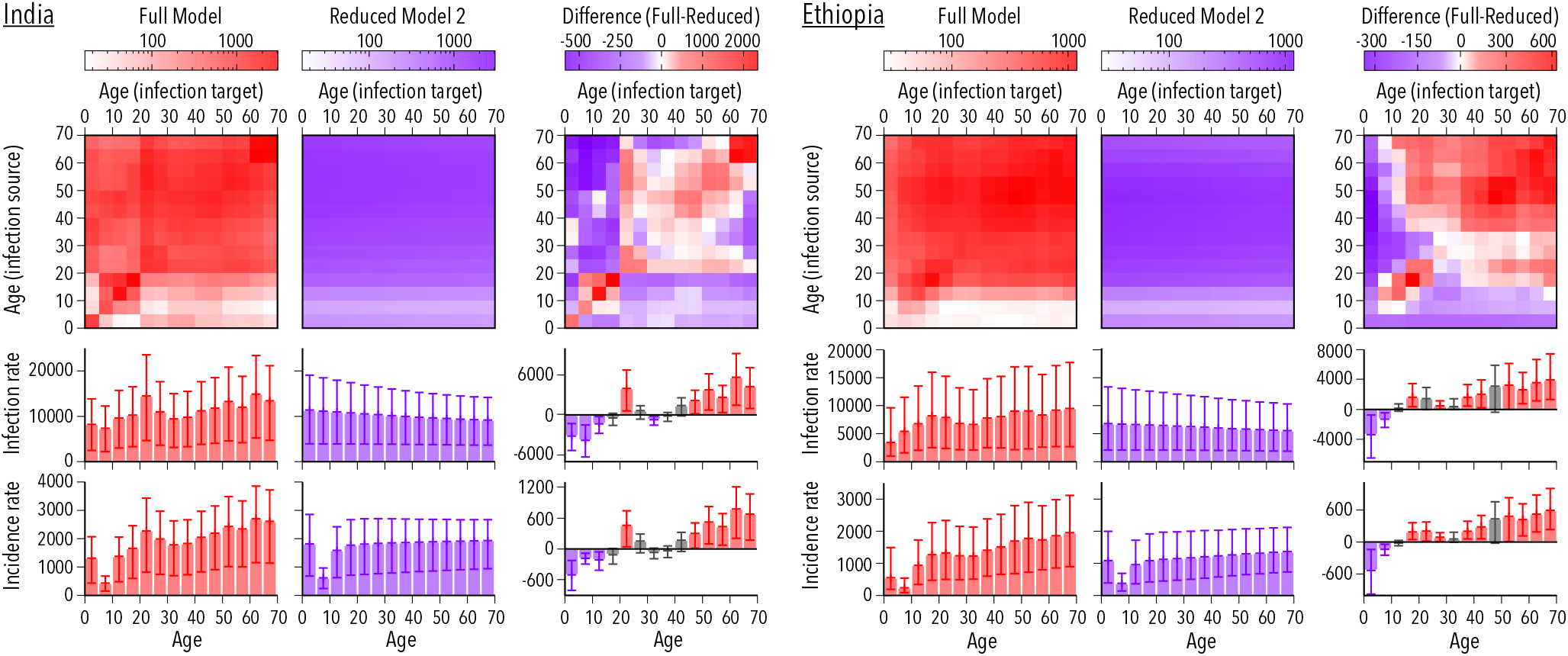
Age-to-age infection rate matrices, and age-specific infection and incidence rates forecasted in 2050 for India and Ethiopia, from full (left) and reduced model 2 (center); with the difference (full - reduced model 2) in the right column in each case. Differences in incidence and infection rates are shown in grey when they are not statistically significant, otherwise they have the color associated to the model for which the rate is higher. When empirical contact patterns are introduced in the model, we observe an increased density of infections close to the diagonal (i.e. contagions taking place between individuals of similar age) and less infections taking place form children to adults or vice-versa. At the level of infection/incidence rates, this translates in an underestimation of burden among adults (and an overestimation among children) associated to assuming contacts homogeneity in the reduced model.

Once we have showed that empirical contact patters adapted from both African and Asian studies produce results that depart significantly from those obtained assuming homogeneous mixing, we interrogated whether the differences between the contact matrices used in both continents (see figure 1C, for example), are significant enough to translate into differences in TB burden forecasts. To do this we conducted an additional test in one country -Ethiopia-, in which we evaluated the differences in the TB burden distribution across ages that emanate from using contact data adapted from African, Asian and, as a control, European studies. The results of this analysis are presented in the SI (figure S9), and evidence that the different contact structures used in this work, derived from different empirical studies, introduce significant differences in the distribution of TB incidence. This emphasizes the importance of the estimation of high quality, country specific data about contact patterns for the production of robust epidemic forecasts in age-structured models.

Finally, we tested whether significant differences regarding age-specific distributions of incident cases can also be observed between the full and the reduced model 2 in a series of alternative modeling scenarios. The results of these tests (analogous to those presented in the SI figures S4-S7 for the effects of demographic dynamics) are gathered in the figure S10 of the SI, and indicate that the effects of empirical contact patterns on TB burden distributions are robustly significant under a wide spectrum of alternative situations.

Summarizing this part, and despite the reduced effect observed on aggregated rates, we showed that including empirical contact structures on TB model dynamics reshapes the transmission patterns among age groups, and generates significant differences in age specific infection and incidence rates. Additionally, we showed that considering different matrices estimated from studies conducted in different geographical areas significantly impact the projected burden distributions.

## Discussion

The model presented here was specifically designed to provide a suitable description of TB transmission dynamics in situations where demography is evolving at the same time that the epidemics unfolds. Importantly, we showed that considering current populations’ aging trends in TB transmission models is followed by a systematic increase in burden forecasts. This worrisome result can be understood in terms of the known mechanisms whereby age affects the transmission dynamics of the disease. In TB, adults are affected by higher age-specific burden levels than children, and, at the same time, they are more efficient spreaders, given their increased tendency to develop infectious forms of pulmonary TB (8). As a consequence, considering populations’ aging translates in higher burden forecasts, simply by increasing the fraction of older individuals with respect to children.

These results suggest that the decay in TB burden levels that has been observed in most countries during the last decades might be harder to sustain than previously anticipated. Under this view, the socio-economic and Public Health improvements that made possible the recent decline of TB world-wide would need to be intensified in many countries if the goal of TB eradication is to be pursued before 2050, at the same time that global aging of human populations unfolds.

Furthermore, our model incorporates a data-driven description of the dependency of TB transmission routes with age, which, we have showed, exerts a significant influence on the forecasted age-distribution of the disease burden, by reshaping infection routes. These results will impact the evaluation and comparison of novel epidemiological interventions, mostly if they are conceived to target specific age-strata. That is the case of new preventive vaccines aimed at substituting or improving BCG, and their application through age-focused vaccination campaigns. In this context, previous works have concluded that a quick immunization of young adults through vaccination campaigns focused on adolescents is expected to produce a faster decline in TB incidence than an alternative strategy based on newborns’ vaccination (21). Our results would further reinforce this hypothesis, to the extent that empirical contact patterns are followed by a relative increase of TB among adults with respect to children. It has to be noted, though, that in the decision of what is the optimal age-group to target in a hypothetical immunization campaign for a new vaccine, at least two additional aspects have to be considered, namely, whether the new vaccine is conceived to boost or substitute BCG, and whether previous exposure to environmental antigens are expected to compromise vaccine performance (via “blocking” (28, 29)). BCG substitutes, and/or vaccines susceptible to lose immunogenicity due to exposure to mycobacterial antigens of individuals before vaccination might not be eligible for adolescent immunization anyways.

Despite all these improvements introduced in this work, our approach is not exempt from the strong limitations that affect all TB transmission models operating at this level of resolution. The outcomes of our model depend on a series of epidemiological parameters and initial burden estimates that are subject to strong sources of uncertainty. Even though we have registered these uncertainties and propagated them to the final model outcomes, future improvements and reassessment of these pieces of input data are generally expected to impact the quantitative outcomes of the model, and to further delimitate the uncertainty ranges here reported. As a first example, international Health Authorities come insisting on the importance of implementing systematic surveys of TB prevalence in many countries, as a means towards more accurate TB burden evaluations. Accordingly, they revise and update their burden estimates on a regular basis, as new data become available, which obviously impacts model calibration and results. Furthermore, the demand of epidemiological studies aimed at obtaining updated estimates of key epidemiological parameters in current epidemiological settings has been pointed out as a primary need for the development of more reliable TB models (30). Similarly, we have seen here the importance of obtaining data of contact patterns specific for each setting, by showing that different contact structures inferred from studies conducted in different parts of the world yield to significantly different distributions of TB burden across age (SI appendix, figure S9). Importantly, the interpretation of these burden distributions of TB across age is hindered by the limited quality of the data available regarding TB distribution across age; which makes adventurous any comparison between model and data. For example, current WHO data-structure only splits TB incidence into two major age groups (0-14 vs 15+), with alleged, heavy under-reporting biases among children.

All these considerations, taken together, evidence the need of further studies, spanning from the implementation of systematic surveys that could unlock more accurate burden estimations (either aggregated or, very importantly, age-specific), to the re-estimation of key epidemiological parameters and contact patterns in specific epidemic settings.

Despite those limitations, in this work we have showed that abandoning the simplifications of constant demography and homogeneous contacts shared by previous models of TB transmission is not just technically feasible, but it has significant effects on model outcomes. Remarkably enough, TB is not the only disease where long characteristic time-scales and strong age-dependencies concur (31, 32), which, despite the specific details of the transmission dynamics of each case, implies that similar corrections to what we have proposed here for the case of TB might be pertinent to correct bias of current epidemic models of other diseases too.

## Materials and Methods

### Tuberculosis Natural History

The description of the Natural History of the disease that we use in our model (Figure 1A) is largely based on previous works by C. Dye and colleagues (8, 20), with lesser variations to make it compatible the structure of data reported by WHO regarding disease type and treatment outcomes (27). Specifically, we deal with a compartmental, age-structured model based on ordinary differential equations, which was implemented in programming language C through a fourth order Runge-Kutta algorithm (time step= 1 day). The model presents two different latency paths to disease –fast and slow– and six different situations of disease, depending on its etiology (non pulmonary, pulmonary smear negative or pulmonary smear positive, characterized by an increasing infectiousness)– and on treatment status. After disease, we explicitly consider the main treatment outcomes included in WHO data schemes: treatment completion, default, failure and death, as well as natural recovery. The natural history model and transitions between the different states, including exogenous reinfections, endogenous reactivations, mother-child transmission and smear progression (i.e. the transition from smear negative to positive during an episode of active TB)(7, 8, 13, 20, 27, 33, 34) are thoroughly detailed in the SI Appendix (see Figure S11 there).

### Age-structure and demographic evolution

The transmission dynamics defined by the Natural History described above is executed in parallel in n=15 age-groups of a span of five years each, except for the last one, which contains all individuals older than 70 (omitted from figures 3-4 to facilitate visual reading of the scales). The internal transitions between disease states within age-groups are then complemented by transitions between age-groups representing individuals’ aging (figure 1B). That defines, per each age group, an a-priori evolution term 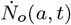 that describes the un-corrected time derivative of the population in age group *a*, at time *t*. Then, to make our demographic structures reproduce the curves reported by the UN Population Division, these empirical data series are fitted to smooth polynomials that are then derived to obtain 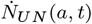 Finally, a correction term 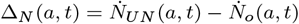, is added to the uncorrected evolution in such a way that the final time derivative of the demographic structure, defined as 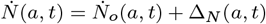 verifies 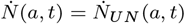 by construction. This, along with the initialization of the population structures according to the UN data, ensures that the evolution of the demography reproduces the UN prospects for all countries and time-points. The correction term Δ_*N*_(*a*, *t*) represents the population variations that occur for causes foreign to TB: new births -introduced as susceptible individuals in the first age group, except for the fraction that undergoes peri-natal infection (see SI section 1.1.10)-, as well as TB-unrelated deaths and migrations, which are distributed, for the rest of the age-groups, among the different disease states proportionally to their respective volumes (see SI section 2.5 for further details).

### Empirical contact patterns

empirical data of age-dependent contact patterns have been adapted from statistical surveys conducted in different countries in Africa, Asia and, as a control, Europe. In each case, contact matrices from studies conducted in different countries of the same continent (Kenya (22), Zimbabwe (23) and Uganda (24) in Africa; China (25) and Japan (26) in Asia and Belgium, Germany, Finland, Great Britain, Italy, Luxembourg, Netherlands and Poland in Europe (14)) have been processed according to the following steps.

First, contact matrices from each study ξ_*s*_(*a*, *a*′) are corrected to preserve symmetry (i.e. to make the total number of contacts between age-groups *a* and *a*′ compatible with survey responses from both groups, conditioned to the demographic structure of the population of each study) and normalized to a common scale. Then, matrices corresponding to studies made on the same continent are averaged, weighted according to the number of participants in each study. As a result, we obtain one matrix per region ξ_*reg*_(*a*, *a*′), which also guarantees that the reports of the contact frequency between *a* and *a*′ are compatible, generating the same number of total contacts, given the demography of the region (i.e. the union of the countries being averaged at the time of the studies):

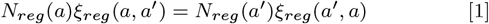

Secondly, to be able to use these averages in specific settings with different demography, we interpret the matrices ξ_*reg*_(*a*, *a*′) as the product of two nuisance factors: the fraction of individuals in *a*′ that exist in the population: 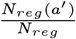, and an auxiliary matrix π_*reg*_(*a*, *a*′):

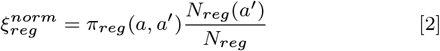

Under this interpretation, the auxiliary matrices π_*reg*_(*a*, *a*′) capture the “intrinsic” intensity of contacts between groups a and a’, once the effect of the demography has been removed, except for a common scale factor.

Next, the matrices π_*reg*_(*a*, *a*′) of each region, as inferred from eq. 2, are adapted to the specific demography of the countries analyzed in this work. Contacts derived from studies conducted in Asia are applied in India and Indonesia, while contacts proceeding from the african studies are applied in Nigeria and Ethiopia. (European contacts are only used as a control in the supplementary appendix). This yields to the following country-specific matrices:

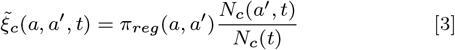

which allow us to incorporate the influence of the evolving demography on the contact structure of our model automatically Finally, ξ_*c*_(*a*, *a*′, *t*) is normalized dynamically at each time step to obtain the final contact matrices used in our model, denoted as ξ_*c*_(*a*, *a*′, *t*). These matrices represent, at any time, the contact frequency that an individual of age a has with individuals of age *a*′, relative to the overall frequency of contacts that any individual has with anyone else in the system (see section 2.2 and 2.3 of the SI, and figure S12 for further details).

### Data flux and model calibration

The flux of data is summarized in figure 1C. The model makes use of four different types of inputs, including: 1. Each of the 19 literature-based epidemiological parameters (7, 8, 13, 20, 33, 34) (SI table S11). 2. TB burden and treatment outcome proportions (reported at the WHO TB database (27), accessed on November 2016, see SI table S12). 3. Contact patterns (estimated from different survey studies conducted in Africa (22–24) and Asia (25, 26)) and 4. Demographic prospects reported in the UN Population Division database (accessed on November 2016).

All these input data are integrated at the step of model calibration, whose goal is to reproduce the time-series of aggregated incidence and mortality reported by the WHO for each country in the period 2000-2015. To achieve this goal, the initial conditions of the system and the values of the only two parameters that do not proceed from bibliographic sources (the scaled infectiousness and the diagnosis rate) are estimated for each country. This procedure is completed using the Levemberg-Marquard optimization algorithm implemented in the C library levmar (see SI section 2.8, figure S13). These two fitted parameters, which define a scale for the number of secondary infections caused by each infectious agent, as well as for the number of cases diagnosed per unit time in each country (SI figure S14), are allowed to vary in time, as in previous works (20), to illustrate socio-economic improvements that might impact the ability of Public Health Systems to better control the disease and restrain its transmission. Finally, the estimates of the initial conditions and the fitted parameters are integrated with the rest of inputs to produce model forecasts up to 2050.

### Uncertainty & Sensitivity Analysis

The uncertainty of each independent source of input data was propagated to model forecasts. The contribution to overall uncertainty assigned to each type of input (epidemiological parameters, WHO estimates of TB burden and treatment outcomes, contact patterns and demographic prospects) has been calculated by repeating model calibration and forecast steps in a series of alternative scenarios where each uncertainty source is shifted sequentially from its expected value to its confidence interval limits. Finally, the deviations from the central estimate that correspond to these alternative scenarios are aggregated assuming mutual independence, and linearly weighted to generate the final confidence intervals showed in figure 2 for aggregated burden projections and in figures 4-5 for incidence and infection rates within age-group. In the SI section 1.3 (figure S3), the individual contribution of all single sources of uncertainty on aggregated incidence and mortality is disclosed, with red (blue) bars representing changes in burden rates associated to an increase (decrease) of the parameters/uncertainty sources from the central values. An important feature of this method is that it allows us to test how sensitive are our forecasts to inputs’ uncertainty upon model re-calibration, instead of testing the intrinsic sensitivity of the dynamics of the non-calibrated model to each input (see SI section 4, for further details).

### Further specifications

For further model details, including definitions of model states, explicit enunciation of model differential equations, parameters values and uncertainties, the reader is referred to sections 2 to 4 of the SI.

## ACKNOWLEDGMENTS

SA was supported by the FPI program of the Government of Aragón, Spain, and JS by the postdoctoral training program for non-residents of Quebec from the Fonds de recherche du Québec - Santé (FRQS) and by the Canadian Institutes of Health Research (CIHR) through a Banting fellowship. This work was partially supported by Gobierno de Aragón/Fondo Social Europeo’, by MINECO and FEDER funds through grants FIS2014-55867-P and BIO2014-52580P, by Project FIS 15/0317 to SS and MJI, by Project TBVAC2020 (643381) funded by the European Commission H2020 to CM and DM, and by the EC Proactive project MULTIPLEX (contract no. 317532) to YM. The funders had no role in study design, data collection and analysis, decision to publish, or preparation of the manuscript. We also thank M. Gutierrez for assistance with figures.

## References

1. Dormandy T (1999) The white death: a history of tuberculosis. (Hambledon Press, London).

2. Borgdorff MW, Floyd K, Broekmans JF (2002) Interventions to reduce tuberculosis mortality and transmission in low-and middle-income countries. Bull World Health Organ 80(3):217–227.

3. Lienhardt C, et al. (2012) Global tuberculosis control: lessons learnt and future prospects. Nat Rev Microbiol 10(6):407–416.

4. World Health Organization (2017) Global tuberculosis report 2017. (Geneva).

5. Dye C, Williams BG (2000) Criteria for the control of drug-resistant tuberculosis. Proc Natl Acad Sci USA 97(14):8180–8185.

6. Boily M, Lowndes C, Alary M (2002) The impact of hiv epidemic phases on the effectiveness of core group interventions: insights from mathematical models. Sex Transm Infect 78 Suppl 1:i78–i90.

7. Korenromp EL, Scano F, Williams BG, Dye C, Nunn P (2003) Effects of human immunodeficiency virus infection on recurrence of tuberculosis after rifampin-based treatment: an analytical review. Clin Infect Dis 37(1):101–12.

8. Abu-Raddad LJ, et al. (2009) Epidemiological benefits of more-effective tuberculosis vaccines, drugs, and diagnostics. Proc Natl Acad Sci USA 106(33):13980–5.

9. Garnett GP, Cousens S, Hallett TB, Steketee R, Walker N (2011) Mathematical models in the evaluation of health programmes. Lancet 378(9790):515–525.

10. Byng-Maddick R, Noursadeghi M (2016) Does tuberculosis threaten our ageing populations? BMC Infect Dis 16(1):119.

11. Lobato MN, Cummings K, Will D, Royce S (1998) Tuberculosis in children and adolescents: California, 1985 to 1995. Pediatr Infect Dis J 17(5):407–411.

12. Dye C, Williams BG (2008) Eliminating human tuberculosis in the twenty-first century. J R Soc Interface 5(23):653–662.

13. Marais B, et al. (2004) The natural history of childhood intra-thoracic tuberculosis: a critical review of literature from the pre-chemotherapy era [state of the art]. Int J Tuber Lung Dis 8(4):392–402.

14. Mossong J, et al. (2008) Social contacts and mixing patterns relevant to the spread of infectious diseases. PLoS Med 5(3):e74.

15. Del Valle SY, Hyman JM, Hethcote HW, Eubank SG (2007) Mixing patterns between age groups in social networks. Soc Networks 29(4):539–554.

16. Melegaro A, Jit M, Gay N, Zagheni E, Edmunds WJ (2011) What types of contacts are important for the spread of infections? using contact survey data to explore European mixing patterns. Epidemics 3(3):143–151.

17. Miller E, et al. (2010) Incidence of 2009 pandemic influenza A H1N1 infection in England: a cross-sectional serological study. Lancet 375(9720):1100–8.

18. Birrell PJ, et al. (2011) Bayesian modeling to unmask and predict influenza A/H1N1pdm dynamics in London. Proc Natl Acad Sci USA 108(45):18238–43.

19. UN (2016) Population division database. http://esa.un.org/unpd/wpp/index.htm (accessed November 2016).

20. Dye C, Garnett GP, Sleeman K, Williams BG (1998) Prospects for worldwide tuberculosis control under the who dots strategy. Lancet 352(9144):1886–91.

21. Knight GM, et al. (2014) Impact and cost-effectiveness of new tuberculosis vaccines in lowand middle-income countries. Proc Natl Acad Sci 111(43):15520–15525.

22. Kiti MC, et al. (2014) Quantifying age-related rates of social contact using diaries in a rural coastal population of Kenya. PloS one 9(8):e104786.

23. Melegaro A, et al. (2017) Social contact structures and time use patterns in the Manicaland province of Zimbabwe. PloS one 12(1):e0170459.

24. le Polain de Waroux O, et al. (2017) Characteristics of human encounters and social mixing patterns relevant to infectious diseases spread by close contact: A survey in southwest Uganda. bioRxiv p. 121665.

25. Read JM, et al. (2014) Social mixing patterns in rural and urban areas of southern China. Proc R Soc Lond B Biol Sci 281(1785):20140268.

26. Ibuka Y, et al. (2015) Social contacts, vaccination decisions and influenza in Japan. J Epidemiol Community Health pp. jech–2015.

27. World Health Organization (2016) Tuberculosis database. http://www.who.int/tb/country/en/index.html (accessed November 2016).

28. Barreto ML, et al. (2014) Causes of variation in BCG vaccine efficacy: examining evidence from the BCG REVAC cluster randomized trial to explore the masking and the blocking hypotheses. Vaccine 32(30):3759–3764.

29. Arregui S, Sanz J, Marinova D, Martín C, Moreno Y (2016) On the impact of masking and blocking hypotheses for measuring the efficacy of new tuberculosis vaccines. PeerJ 4:e1513.

30. Dowdy DW, Dye C, Cohen T (2013) Data needs for evidence-based decisions: a tuberculosis modeler’s ‘wish list’. Int J Tuberc Lung Dis 17(7):866–77.

31. Hontelez JA, et al. (2011) Ageing with HIV in South Africa. AIDS (London, England) 25(13).

32. Griffin JT, Ferguson NM, Ghani AC (2014) Estimates of the changing age-burden of plasmodium falciparum malaria disease in sub-Saharan Africa. Nat Commun 5.

33. Picon PD, et al. (2007) Risk factors for recurrence of tuberculosis. J Bras Pneum 33(5):572–8.

34. Pillay T, Khan M, Moodley J, Adhikari M, Coovadia H (2004) Perinatal tuberculosis and HIV-1: considerations for resource-limited settings. Lancet Inf Dis 4(3):155–65.

